# Reversal Learning Phenotypes are Linked with Novel Genetic Loci in Diversity Outbred Mice

**DOI:** 10.1101/2022.01.29.478259

**Authors:** Jared R. Bagley, Lauren S. Bailey, Leona H. Gagnon, Hao He, Vivek M. Philip, Laura G. Reinholdt, Lisa M. Tarantino, Elissa J. Chesler, James D. Jentsch

**Author notes:** Corresponding Author: Jared Bagley, Department of Psychology, State University of New York at Binghamton Binghamton NY, 13902.

## Abstract

Impulsive behavior and impulsivity are heritable phenotypes that are strongly associated with risk for substance use disorders in human subjects. Consequently, identifying the neurogenetic mechanisms that influence impulsivity may also reveal novel biological insights into addiction vulnerability. Past studies from our laboratory using the BXD and Collaborative Cross (CC) recombinant inbred mouse panels have revealed that behavioral indicators of impulsivity measured in a reversal learning task are heritable and are genetically correlated with aspects of intravenous cocaine self-administration. Genome wide linkage studies in the BXD panel revealed a quantitative trait locus (QTL) on chromosome 10, but the specific genes affecting this trait remain elusive. To achieve greater precision in our mapping efforts, we have turned to Diversity Outbred (DO) mice. A total of 392 DO mice (230 males, 295 females) were successfully phenotyped using the same reversal learning test utilized in our earlier studies. Our primary indicator of impulsive responding, a measure that isolates the relative difficulty mice have with reaching performance criteria under reversal conditions, revealed a genome wide significant QTL on chromosome 7 (max LOD score = 8.73, p<0.05). A measure of premature responding akin to that implemented in the 5-choice serial reaction time task yielded a suggestive QTL on chromosome 17 (max LOD score = 9.14, p<0.1). Positional candidate genes were prioritized (*2900076A07Rik, Wdr73 and Zscan2)* based upon expression QTL data we collected in DO and CC mice and analyses using publicly available gene expression and phenotype databases. These findings may advance understanding of the genetics that drive impulsive behavior and enhance risk for substance use disorders.

---

Many people initiate experience with potentially addictive substances, yet only a fraction of those develop a clinically impairing substance use disorder (Wagner & Anthony, 2002). Stimulant drugs, including cocaine, are no exception; a majority of people who initiate cocaine use in their lifetime will not become addicted to it (Wagner & Anthony, 2002). The transition from subclinical, recreational use to a SUD is influenced by both genetic and environmental factors, as well as interactions between them (Goldman et al., 2005; Palmer et al., 2009; Prom-Wormley et al., 2017); at least 50% of the risk for developing a cocaine use disorder is attributable to genetic variation (Goldman et al., 2005). Moreover, genetic risk for cocaine addiction is, to a substantial degree, shared with other illicit drugs of abuse (Dick, 2016; Goldman et al., 2005; Prom-Wormley et al., 2017), meaning that identifying genetic loci regulating cocaine-related behaviors indirectly informs us about the genetics that influence clinically-impairing use of other substances. To date, the specific genes and gene networks that influence the vulnerability to transition to compulsive drug-seeking and -taking remain mostly unknown. This knowledge gap represents a barrier the limits the ability to design and develop effective prevention and treatment options.

Impulsivity, which can be described as either difficulty with inhibiting impulsive reward pursuit or consumption (impulsive action) and/or as impulsive reasoning about reward-related behaviors (impulsive choice) (Dalley et al., 2011; J. D. Jentsch et al., 2014; Winstanley et al., 2010), has been repeatedly linked with the initiation of drug and alcohol use and progression into an SUD (Cervantes et al., 2013a; Dalley et al., 2007; J. D. Jentsch et al., 2014; Winstanley et al., 2010). Although impulsive action and choice phenotypes may be distinct in terms of underlying biological mechanisms (Broos et al., 2012; Dalley et al., 2008; Dalley & Robbins, 2017; J. D. Jentsch et al., 2014; MacKillop et al., 2016), both predict aspects of the response to cocaine in animal models and humans. For example, inter-individual differences in impulsivity predict the propensity to: 1) experience altered subjective effects of potentially addictive substances (Weafer & De Wit, 2013); 2) initiate cocaine intravenous self-administration (IVSA) (Anker et al., 2009; Belin et al., 2008; Cervantes et al., 2013a; Dalley et al., 2007; Perry et al., 2005, 2008); 3) transition to habitual/inflexible use (Belin et al., 2008; Broos et al., 2012); and 4) relapse after periods of withdrawal or abstinence (Adinoff et al., 2016; Broos et al., 2012; Perry et al., 2008). Our work has revealed that the predictive relationship between impulsive action and cocaine IVSA is attributable to a genetic correlation, also known as co-heritability (Cervantes et al., 2013a).

Impaired impulsive action may result from deficient inhibitory control over behavior and ultimately manifest as a proclivity to persist in drug use despite negative outcomes. Laboratory tasks that measure inhibitory control provide opportunities to investigate the biology of behavioral flexibility, including indirectly uncovering the neurogenetic mechanisms of addiction vulnerability. One procedure, called reversal learning, measures a subject’s ability to suppress the response to a previously reinforced behavior when response-outcome contingencies unexpectedly change (Izquierdo & Jentsch, 2012). Reversal learning deficits are associated with drug use and SUDs, both in laboratory animals and human subjects, and therefore may be informative of biological factors that drive impulsivity and subsequent risk for SUDs (Calu et al., 2007; Camchong et al., 2011; Cervantes et al., 2013a; Gullo et al., 2010; Izquierdo & Jentsch, 2012; J. Jentsch, 2002; Smith et al., 2015).

Reversal learning is influenced by genetic variation in rodent populations that can be utilized to map associated genetic loci (Bailey et al., 2021; Laughlin et al., 2011). Laboratory rodent populations offer some distinct advantages in forward genetic approaches. Genetically diverse populations can be tested in prospective, highly controlled experimental designs that can reveal quantitative trait loci (QTL) associated with impulsive traits and addiction liability. Concurrent study of genome-wide transcript expression can support discovery of candidate genes and gene networks that affect behavioral flexibility.

The Diversity Outbred (DO) mice and Collaborative Cross (CC) inbred strains populations were developed by interbreeding a highly genetically diverse set of founder strains (Chesler, 2014; Churchill et al., 2004, 2012; Philip et al., 2011; Threadgill & Churchill, 2012). High genetic diversity can expand phenotypic distributions and provide unique opportunities for discovery of variants that drive extreme phenotypes (Chesler, 2014). Reversal learning is heritable in CC strains and their founders (Bailey et al., 2021), indicating these populations may be suitable for genetic dissection of this trait. The DO mice may thus be utilized for relatively high-resolution QTL mapping studies. the CC strains support discovery of genetic correlations among gene expression and behavioral traits, in a fully replicable population that allows for cumulative research and inter-study analyses.

Here, we describe QTL mapping for reversal learning using DO mice. We also advance positional candidate discovery using reversal learning data from the CC strains along with complementary whole-transcriptome gene expression measures generated from bulk RNA sequencing of striatal tissue (previously described (Bailey et al., 2021; Saul et al., 2020) to advance positional candidate discovery. The striatum is a key brain region of interest in reversal learning performance and SUDs (Bergstrom et al., 2020; Clarke et al., 2008; Cools et al., 2009; Everitt & Robbins, 2013). Collectively, these experiments may reveal genes that moderate reversal learning and enhance understanding of SUD neurogenetics.

## Methods

### Subjects

Diversity outbred (DO) mice (n = 525) and CC strains (n = 33) (Bailey et al., 2021) were born at the Jackson Laboratory, Bar Harbor, ME and maintained there in dedicated mouse colony rooms on a 12:12 h light:dark cycle and at an average temperature of 69–70°F. Food (Lab Diet 5001, ScottPharma Solutions) and water was available *ad libitum* prior to initiation of food restriction and behavioral testing (described below). A nestlet and a disposable dome-shaped shack were provided in the home cage (Shepherd Specialty Papers, Inc., Watertown, TN, USA). Mice were group housed post weaning, transitioned to single housing at 6 weeks of age and maintained under single housing for the duration of testing. All DO/CC mice were tested at JAX by the Behavioral Phenotyping Core, a component of the Systems Neurogenetics of Addiction. Animal studies were performed according to the “Guide for the Care and Use of Laboratory Animals” (National Research Council, 2011) in the AAALAC accredited programs at JAX, after approval by the relevant Institutional Animal Care and Use Committee.

### Novelty-Related Behavioral Testing

The DO mice utilized for reversal learning were initially tested (7-8 weeks of age) for locomotor and novelty related behaviors beginning at 8 weeks of age, as previously described (Saul et al., 2020). These tests included the open field, light-dark box, hole board and a measure of novel place preference. All mice experienced all forms of testing under equivalent protocols and conditions. The data from these studies are not reported here.

### Food Restriction

Prior to the initiation of the reversal learning protocol described below, mice were introduced to a schedule of limited access to chow. Mice were weighed daily during food restriction and percent of free-feeding body weight was calculated by dividing the current weight by the pre-restriction weight. During the limited access to food period, mice were fed once a day; chow quantity provided per day was titrated until mice reach 80%–90% of their pre-restriction weights. Once mice reached their target weights, operant testing began. If, at any point during the testing period, a mouse dropped below 80% of its free feeding weight, the quantity of chow provided was increased. If increased food availability did not lead to a recovery of body weight to ≥80% within a day, it was temporarily returned to *ad libitum* food access until its weight had recovered.

### Reversal Learning

Reversal learning testing began at 9-13 weeks of age. Testing took place in 8.5″ L × 7″ W × 5″ H (21.6 × 17.8 × 12.7 cm) operant conditioning modular chambers (Model ENV-307W, Med Associates Inc.) that were fitted with stainless-steel grid floors (Model ENV-307W-GFW, Med Associates Inc.) and located in sound attenuating cubicles. The operant box contains a horizontal array of five nose poke apertures on one side of the box, and a central food magazine. A house light and white noise maker were positioned within the cubicle above the operant box, as well.

Immediately prior to testing, mice were removed from their home cage by grasping the tail with large, padded forceps and placed inside the operant box. Each mouse was sequentially tested in a series of programs; mice transitioned from program to program individually, as they met criterion performance (see below). Mice underwent the following programs:

Stage 1: *Box habituation*. House light and white noise were active. No reinforcements were provided. Box habituation comprised of one session that lasted 1-h.

Stage 2: *Magazine training*. House light and white noise were active for the duration of the test. During this test, 20–21 μl Original Strawberry Boost (Nestlé HealthCare Nutrition, Inc., Florham Park, NJ) was dispensed into the food magazine every 30 s. The session ended after 1-h or after the mouse received and retrieved 50 rewards, whichever came first. A mouse progressed to Stage 3 when it earned 30 or more rewards within a session.

Stage 3: *Initial operant* (*nose-poke*) *conditioning*. Sessions began with illumination of the house light and activation of the white noise generator; 10-s later, nose poke aperture 3 of 5 (center aperture) was illuminated. A behavioral response that broke the photocell in the aperture (usually, a nose poke) resulted in the extinction of the internal light; in addition, if the beam was broken for a continuous pre-set period of at least 1 (beam break with no additional hold time), 100, or 200 ms (the time requirement varied randomly from trial to trial), the action was reinforced by the delivery of 20–21 μl of Boost solution; after each reinforcer was retrieved, a new trial was initiated 1.5-s later (signaled by illumination of the center nose poke aperture). If a response was initiated but not sustained for the preset period, a time out period of 2-s occurred, during which time the central nose poke light and house light were extinguished. If a mouse did not voluntarily respond in the center hole for at least 15 minutes, that hole was baited with a Boost-saturated cotton swab. Daily sessions lasted up to 1-h but were terminated prior to that time if an individual mouse earned 50 reinforcers. Each mouse was tested daily on this stage until it received at least 50 reinforcers in a single session, at which time it progressed to the next stage.

Stage 4: Mice were tested under the same basic conditions outlined in Stage 3, except that a minimum duration nose poke of 100- or 200-ms was required to produce reinforcement. If a mouse had not responded in the central illuminated hole for 15 min, that hole was again baited with a Boost-saturated cotton swab. When the mouse earned 50 reinforcers in a single session, it progressed to Stage 5. If the mouse had not met criteria after 10 days, it was regressed to Stage 3. If the subject returned to Stage 4 but did not meet criteria after another 10 test days, it was removed from the study because of failure to progress. Across all Stages, a mouse could only regress once. For example, if a mouse did not pass Stage 4 in 10 days and regressed to Stage 3, then later did not pass Stage 5 within 10 days, the mouse was removed from the study.

Stage 5: In this phase, mice were tested under the same basic conditions as outlined in Stages 3 and 4, except that a minimum duration nose poke of 100-, 200-, or 300-ms was required to trigger reinforcement delivery. If a mouse did not respond in the center illuminated hole for 15 minutes, that hole was baited with a Boost-saturated cotton swab. When the mouse earned 50 reinforcers in a single session, it progressed to the *Discrimination learning* stage. If the mouse did not meet passing criteria after 10 days, they regressed to Stage 4. If the subject returned to Stage 5 but still did not meet criteria after another second of 10 test days, it was removed from the study because of regression failure.

Stage 6: *Discrimination learning*. As above, session onset was signaled by illumination of the house light and activation of the white noise generator; trial onset was signaled by illumination of the center nose poke aperture. As in Stage 5, mice were required to first complete an observing response into the central aperture of 100-, or 200-ms duration; any nosepokes into the target (flanking) holes before completing the observing response and successfully initiating a trial were counted as premature/anticipatory responses. Once a trial was successfully initiated with an observing response, the two apertures flanking the central hole (hole 2 and 4) were illuminated. A response into one of the two apertures (pseudorandomly assigned across strains) resulted in the delivery of a Boost reinforcer (this was counted as a correct choice). Poking into the other hole - or not making any response within 30-s, triggered a time out, during which time the house light was extinguished; these outcomes were counted as an incorrect choice or an omission, respectively. Daily sessions of 1-hr were conducted until learning criteria were met; these criteria included a mouse completing at least 20 trials in a single session and achieving at least 80% accuracy over a running window that included the last 20 trials. A mouse regressed to Stage 5 if it did not complete at least 10 trials for three consecutive days. If 300 trials were completed without meeting passing criteria, the mouse was removed from the study because of Stage 6 failure.

Stage 7: *Reversal learning stage*. Testing was nearly identical to that described above in Stage 6, with the exception that the reinforcement contingencies associated with the two holes were switched. Testing progressed in daily sessions until animals once again met the same learning criteria rule described above, and the same dependent variables were collected (see below). After reversal was completed, mice were gradually adjusted back onto *ad libitum* feeding. Subjects failed Stage 7 and were removed from the study if 400 trials were completed, or 8 weeks of testing passed, without meeting criteria.

Key dependent variables for the discrimination learning and reversal learning stages were total trials required to reach criteria (TTC) in each stage and premature responding. TTC was calculated as the total number of completed trials (all trials ending in an incorrect or correct response) until it met the performance criteria. The difference in TTC in the reversal stage to TTC in the discrimination learning stage revealed each animal's ability to alter responding under a changing reward contingency, with a non-zero, positive difference score indicating some degree of difficulty with altering its behavior and/or inhibiting the initially trained response.

Premature responses were nose pokes into one of the flanking target holes before a trial is successfully initiated, a measure roughly analogous to that collected in the 5-choice serial reaction time (Bari et al., 2008). Premature responses were separately counted for the correct and incorrect target aperture. Premature responding thus had four values: premature responding in the correct hole at acquisition, premature responding in the incorrect hole at acquisition, premature responding in the correct hole at reversal, and premature responding in the incorrect hole at reversal. All premature responding values were further divided by the animal's TTC in that stage to estimate the average number of premature responses made per trial. Of particular interest is premature responding in the correct hole during acquisition and in the incorrect hole at reversal, as these are the dominant types of responses made.

Other variables measured were the frequency of omissions (total omissions/TTC for each stage); the average proportion of correct trials (total correct trials/TTC; the average trial initiation latency (total trial initiation latency/TTC), which is calculated as the average amount of time that passes between the end of one trial and the successful initiation of the next one; and average reward retrieval time (total reward retrieval time/total correct trials), which is defined as the average amount of time that passes between a reward being administered and the animal’s head entering the magazine.

Key variables of interest were assessed with descriptive statistics (range, mean, standard deviation), and total trials to criterion was assessed by analyses of variance (ANOVA) to determine effects of stage (acquisition and reversal) and sex.

### Genotyping

Tails were removed from each animal at euthanasia, placed into 1.5 mL Eppendorf tubes, and stored in saline at −80°C until DNA extraction. Tail samples were shipped to GeneSeek (Neogen Inc., Lincoln, NE, USA) for DNA extraction and genotyping on the GigaMUGA (N = 500) Illumina array platforms. The GigaMUGA assays 143,259 genetic markers spanning the 19 autosomes and X chromosome of the mouse, with a mean spacing of 18 Kb (Morgan et al., 2015). Markers were optimized for information content in DO mice. Genotypes were imputed to a 69K grid to allow for equal representation across the genome.

### SNP-Based Heritability

Heritability was estimated in DO mice using 112,470 SNP sets after quality control. We first used these SNP sets to construct genetic relationship matrices (GRMs) in DO mice using GCTA (Yang et al., 2011). We then used the restricted maximum likelihood (REML) approach within GCTA on the GRMs, sex as a covariate, to calculate the SNP-based heritability for each reversal learning phenotype.

### Quantitative trait locus mapping

DO genome reconstruction, sample and marker quality control and QTL mapping were carried out using R/qtl2 software (v 0.28) as described previously (Broman, 2014; Broman et al., 2019; Church et al., 2015; Gatti et al., 2014; Svenson et al., 2012). Briefly, R/qtl2 software constitutes a set of functions designed for QTL mapping in multi-parent populations derived from more than two founder strains. R/qtl2 allows users to perform genome scans using a linear mixed model to account for population structure and permit the imputation of SNPs based on founder strain genomes. Sex and generation were included as covariates for association and linkage mapping.

### Linkage mapping

For linkage mapping, we used an additive haplotype model with kinship correction to estimate founder effects for each QTL. We accounted for genetic relatedness between mice by using a kinship matrix based on the leave-one-chromosome-out (LOCO) method (Cheng & Palmer, 2013). The LOCO method was chosen because kinship calculations that include the causative marker are known to produce overly conservative mapping results (King & Long, 2017; Yang et al., 2014). The genome-wide significance thresholds corresponding to *p*-values < 0.01, 0.05, 0.10 and 0.63, for each trait, were calculated using 1000 permutations to create a null distribution of LOD scores. A QTL was deemed significant if the genome-wide *p*-value was less than 0.10, otherwise it was deemed suggestive. When a QTL peak was identified above any of the above thresholds, a 1.5 LOD drop was used to determine the corresponding QTL region (Broman et al., 2019; Gatti et al., 2014).

### Local Association mapping

For each significant and/or suggestive QTL region, we imputed all high-quality SNPs from the Sanger Mouse Genome Project (build REL 1505; (Keane et al., 2011) onto DO genomes and fit an additive genetic model at each SNP. This approach is widely used in human GWAS and increases power and precision by measuring the effects at individual variants by mapping at the two-state SNP level (Gatti et al., 2014).

### Gene Expression

RNA sequencing was performed on striatal tissue collected from 33 CC strains and 369 DO mice (drug naïve), as previously described (Saul et al., 2020). Each strain was tested under a sensitization protocol following exposure to either cocaine or saline control (two groups of mice per strain) as described in Schoenrock et al, 2020. Tissue was collected 24 to 48 hours after the final injection.

### Expression QTL mapping

Briefly, gene expression counts were obtained by summing expected counts over all transcripts for a given gene. eQTL mapping was performed on regression residuals of 17,248 genes using the R/qtl2 package with the founder haplotype regression method. Kinship matrices to correct for population structure were computed with the LOCO method for kinship correction (Gatti et al 2014; http://kbroman.org/qtl2). Sex and generation were included as additive covariates. We then used the interactive, web-based analysis tool QTL viewer (http://34.74.187.222/) to visualize the expression data with profile, correlation, LOD, effect, mediation and SNP association plots. Detailed information about the structure of the QTL viewer objects are available at: https://github.com/churchill-lab/qtl-viewer/blob/master/docs/QTLViewerDataStructures.md.

### Positional candidate gene prioritization

Gene expression and reversal learning data obtained from CC strains (Bailey et al., 2021; Saul et al., 2020) was utilized to prioritize positional candidate genes for the behavioral QTL detected in DO mice. Pearson’s correlations were calculated for strain-level gene expression, in cocaine and saline exposed mice, to reversal learning in the same strains. The reversal difference score and total trials to acquisition and reversal were assessed. Genes with correlations of FDR < 0.25 were considered prioritized candidates.

These candidates were further assessed for genetic association to other traits of potential interest by use of the ePHeWAS tool available on http://systems-genetics.org, which calculates correlations of strain-level gene expression from publicly available databases to all traits in the phenome database on http://genenetwork.org (Mulligan et al., 2017). The striatum and frontal cortex (FC) were selected as regions of interest for this analysis (Bergstrom et al., 2020; Clarke et al., 2008; Cools et al., 2009; Everitt & Robbins, 2013; Goldstein & Volkow, 2002; Hornak et al., 2004; Wise & Robble, 2020). Multiple comparisons were corrected by Bonferroni adjustment.

## Results

### Reversal Learning

DO mice displayed a wide range of performance in reversal learning. During acquisition, total trials to criterion ranged from 20 to 298, with a mean of 81.6 and a standard deviation of 53.9. During the reversal stage, totals trials to criterion ranged from 20 to 400, with a mean of 142.2 and a standard deviation of 73.0. A mixed ANOVA, with stage as a repeated measure and sex as a between-subjects factor revealed main effects of stage [F(1,390)=229.0, p<0.001] (Fig 1A) and sex [F(1,390)=7.8, p=0.005], with males requiring a larger number of trials to reach the preset performance criterion at both stages (male mean ± SEM = 120.8 ± 4.5; female mean ± SEM = 106.0 ± 3.0). A Pearson’s correlation analysis performance on acquisition and reversal data from individual mice revealed a modest correlation (r=0.29, r^2^=0.08, p<0.001) (Fig 1C).

**Figure 1.**
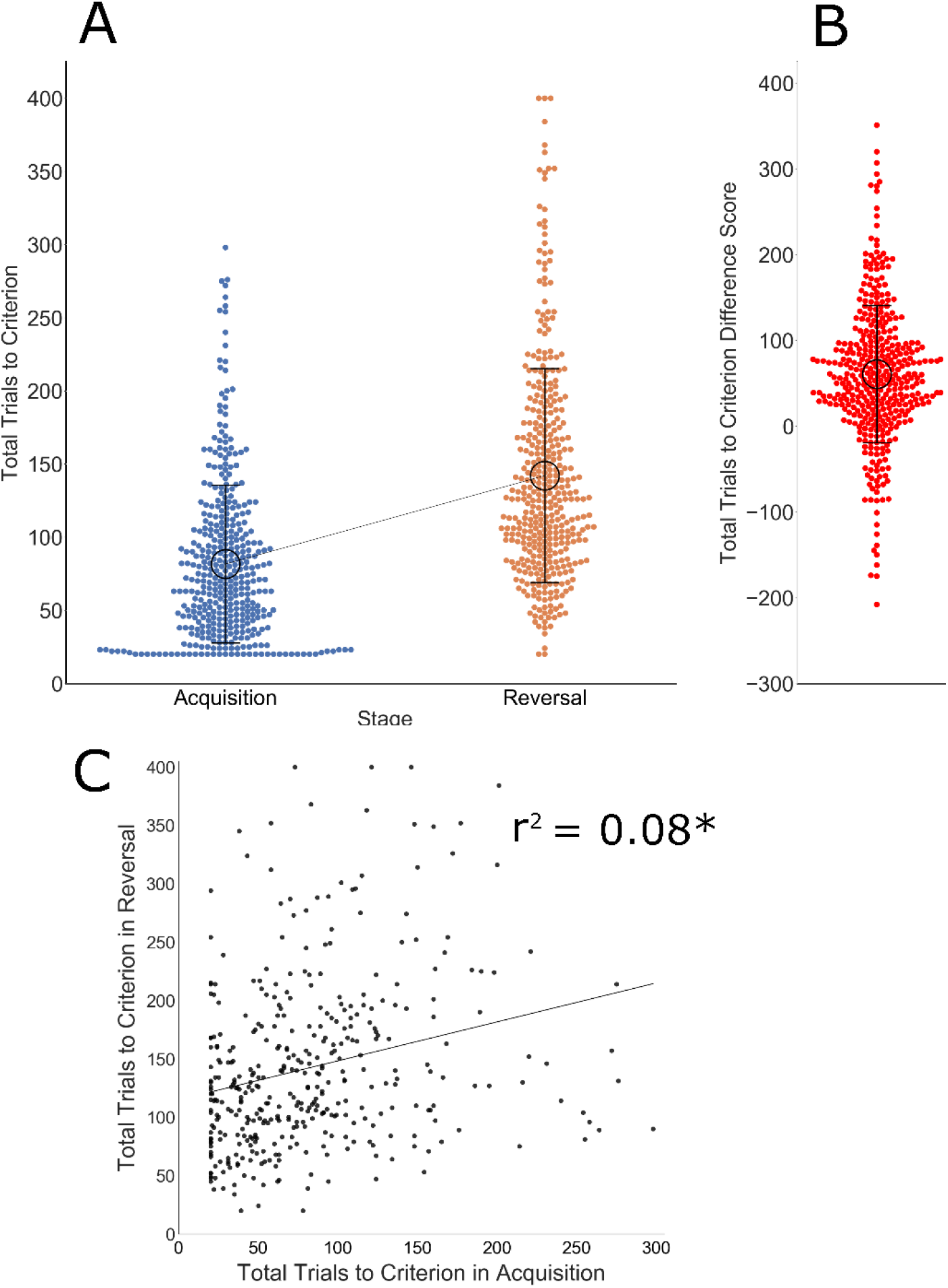
Reversal learning in DO mice. A) As expected, the average number of trials required to reach preset performance criteria were larger in the reversal, as compared to acquisition, stage. DO mice required a wide range of total trials in both the acquisition and reversal learning stages. B) A difference score captures relative difficulty in reaching criterion in the reversal stage. Again, DO mice displayed a broad range of performance and this measure was found to be heritable. C) A significant correlation was detected between acquisition and reversal stages; however, only 8% of variance is shared between these measures.

The difference score (total trials in reversal minus total trials in acquisition) ranged from −208 to 351, with a mean of 60.6, a standard deviation of 80.1 and heritability of 0.06. The DO mean was higher that of CC and founder mice (Bailey et al., 2021); however, variance is similar between the populations (−271 to 383, mean = 37.2, SD = 85.1).

DO mice displayed a wide range of premature responding phenotypes in the correct aperture during the acquisition stage (0 to 8.05 premature responses/trial, mean=0.72, SD=0.82) or in the incorrect aperture during the reversal stage (0.03 to 7.63 premature responses/trial, mean = 1.12, SD = 0.88). The range, mean and variance were greater relative to CC/Founder mice in acquisition (0 to 5.7, mean 0.65, SD = 0.71) and reversal (0 to 5.3, mean = 1.0, SD = 0.80) (Bailey et al., 2021) (Fig. 2). See Table 1 for descriptive statistics of additional variables collected during testing.

**Figure 2.**
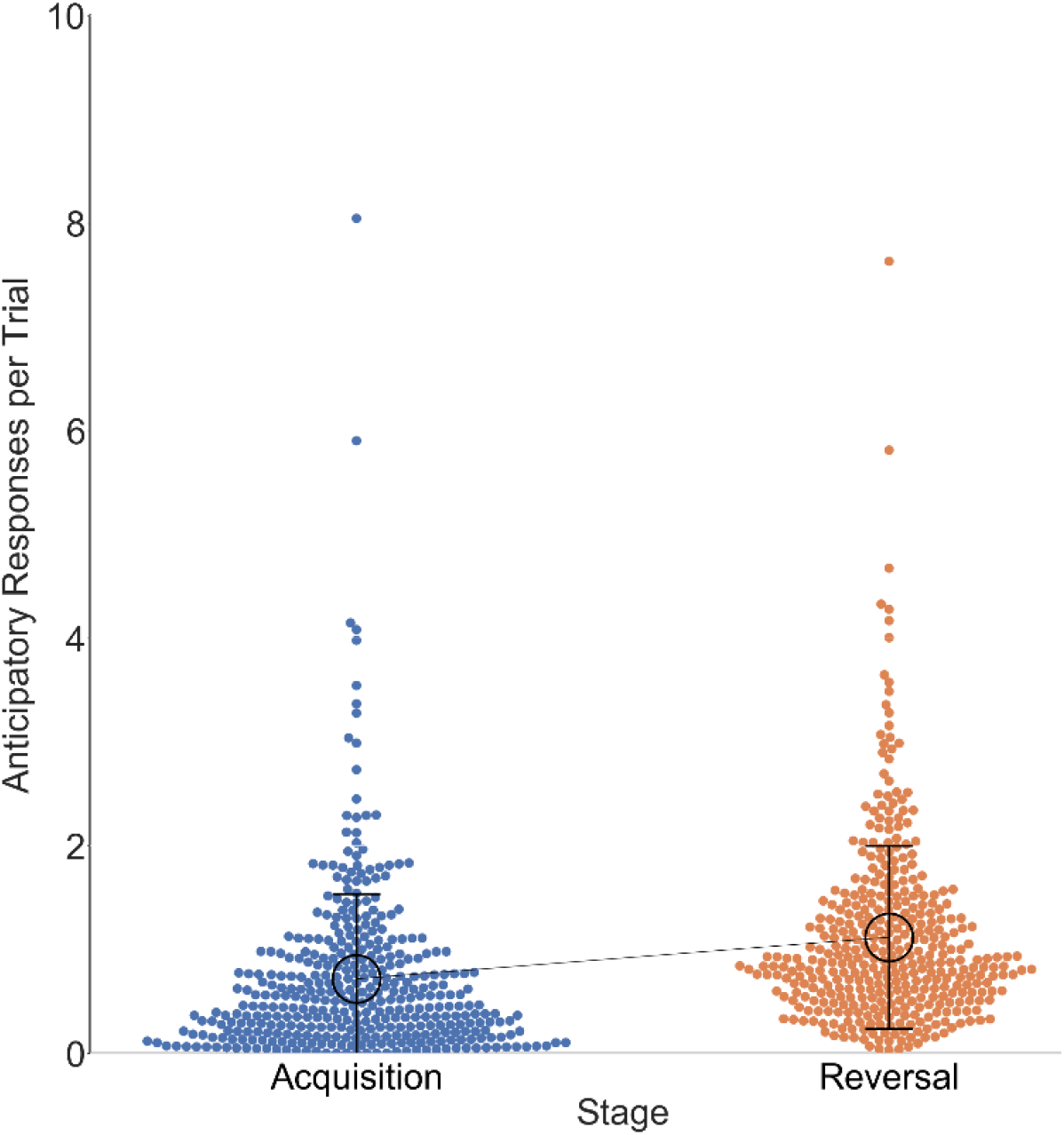
Premature responding in acquisition (correct aperture) and reversal (incorrect aperture) are expressed as a fraction of the total trials initiated. DO mice displayed a broad range of responding in these measures.

**Table 1.** Reversal learning statistics for all DO mice.

| Phenotype | Acquisition |  |  |  |  | Reversal |  |  |  |  |
| --- | --- | --- | --- | --- | --- | --- | --- | --- | --- | --- |
|  | Min | Max | Mean | St. Dev. | Heritability | Min | Max | Mean | St. Dev. | Heritability |
| Days To Criteria | 1 | 20 | 3 | 3 | 0.19 | 1 | 38 | 5 | 5 | 0.03 |
| Total Trials | 20.0 | 298.0 | 81.6 | 53.9 | 0.19 | 20.0 | 400.0 | 142.0 | 73.0 | 0.12 |
| Percent Correct | 35.3 | 95.0 | 62.1 | 12.1 | 0.23 | 20.3 | 80.0 | 47.7 | 10.1 | 0.05 |
| Difference Score | n/a | n/a | n/a | n/a | n/a | -208.0 | 351.0 | 60.6 | 80.1 | 0.06 |
| Premature Correct Responses (per trial) | 0.0 | 8.0 | 0.7 | 0.8 | 0.22 | 0.0 | 6.6 | 0.9 | 0.8 | 0.12 |
| Premature Incorrect Responses (per trial) | 0.0 | 5.2 | 0.5 | 0.5 | 0.24 | 0.0 | 7.6 | 1.1 | 0.9 | 0.33 |
| Total Time (min) | 6.6 | 1195.6 | 153.0 | 181.7 | 0.19 | 14.4 | 2264.6 | 290.0 | 326.1 | 0.04 |
| Trial Initiation Latency (sec., per trial) | 2.0 | 257.8 | 34.7 | 30.8 | 0.18 | 3.3 | 314.6 | 51.9 | 51.2 | 0.14 |
| Reward Retrieval Time (sec., per correct trial) | 7.3 | 58.4 | 19.4 | 6.6 | 0.28 | 8.8 | 54.7 | 21.0 | 7.4 | 0.32 |

A Pearson’s correlation was calculated between the reversal learning difference score and premature responding on the incorrect aperture during the reversal stage. A modest correlation was detected (r = −0.12, p = 0.02, r^2^ = 0.01) (Fig. 3), indicating a large proportion of unshared variance and suggesting these measures may capture distinct phenotypes.

**Figure 3.**
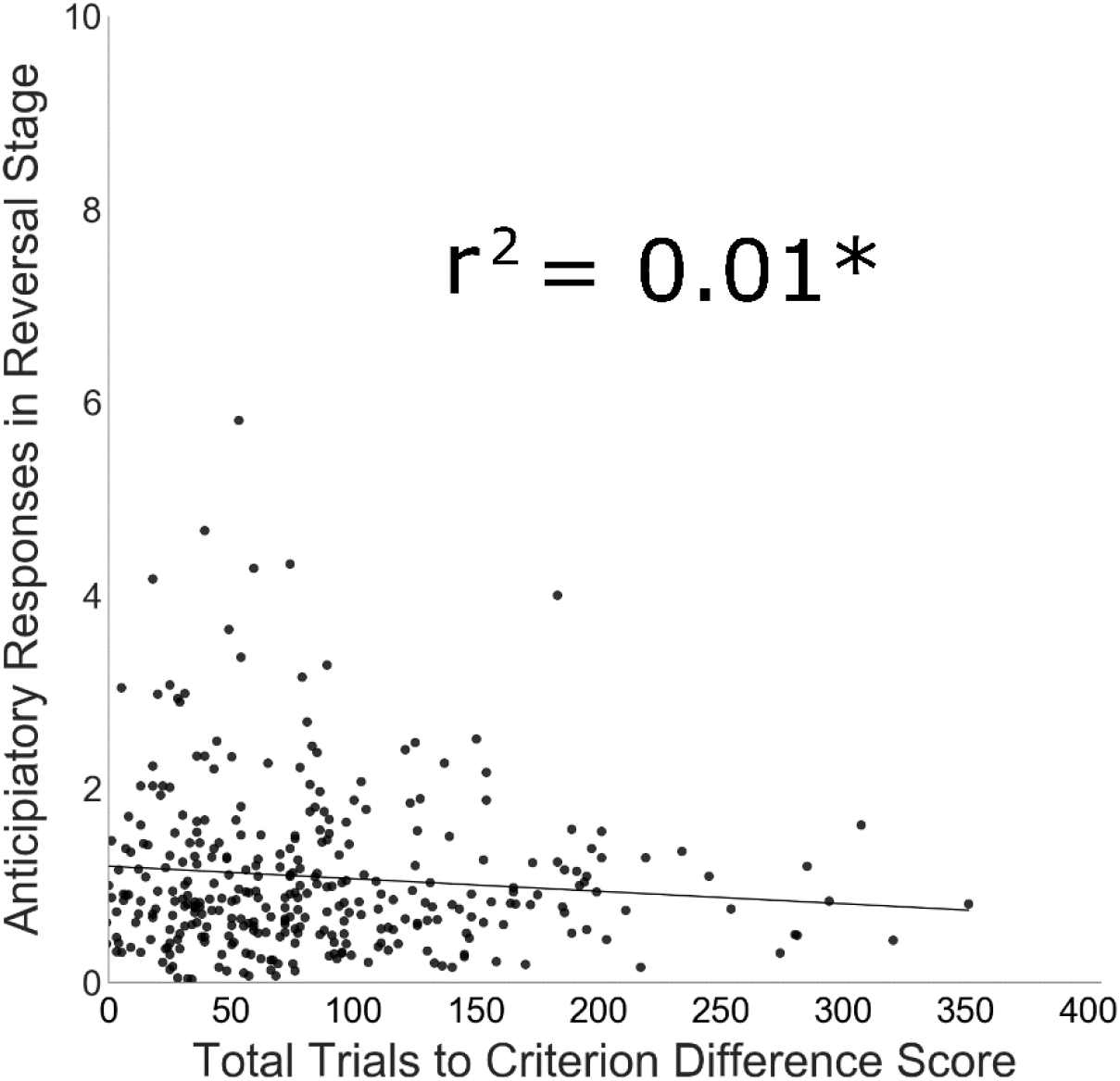
A significant correlation was detected between premature responding in reversal and the reversal learning difference score; however, only 1% of variance is shared, indicating these two measures may capture largely distinct traits. A similar, strain-level r^2^ value (r^2^=0.2, p=0.06) was found for CC strains (Bailey et al. 2021), indicating a similarly small genetic correlation between these traits.

Of the mice that initiated testing, 25% failed to successfully complete reversal learning due either to testing criteria failure (17.2%), health problems (5.1%), technical error (2.1%) or another reason (0.6%). 55.6% of mice that failed were male, suggesting a potential sex-bias in attrition (44.0% of total mice tested were male).

### QTL Mapping

The reversal learning difference score was subject to QTL mapping. A significant QTL on chromosome 7 (position is in GRCm38, Mbp): Chr07, Peak = 80.80581, LOD = 8.725234, Confidence Interval = 80.26511-81.51397) was detected, suggesting a variant(s) at this locus associated with reversal learning performance (Fig. 4A). The additive effects of haplotypes indicated the NZO/HILtJ haplotype associated with positive difference scores (relatively poor reversal learning) and the 129/SvlmJ haplotype associated with negative differences scores (relatively good reversal learning) (Fig 4B).

**Figure 4.**
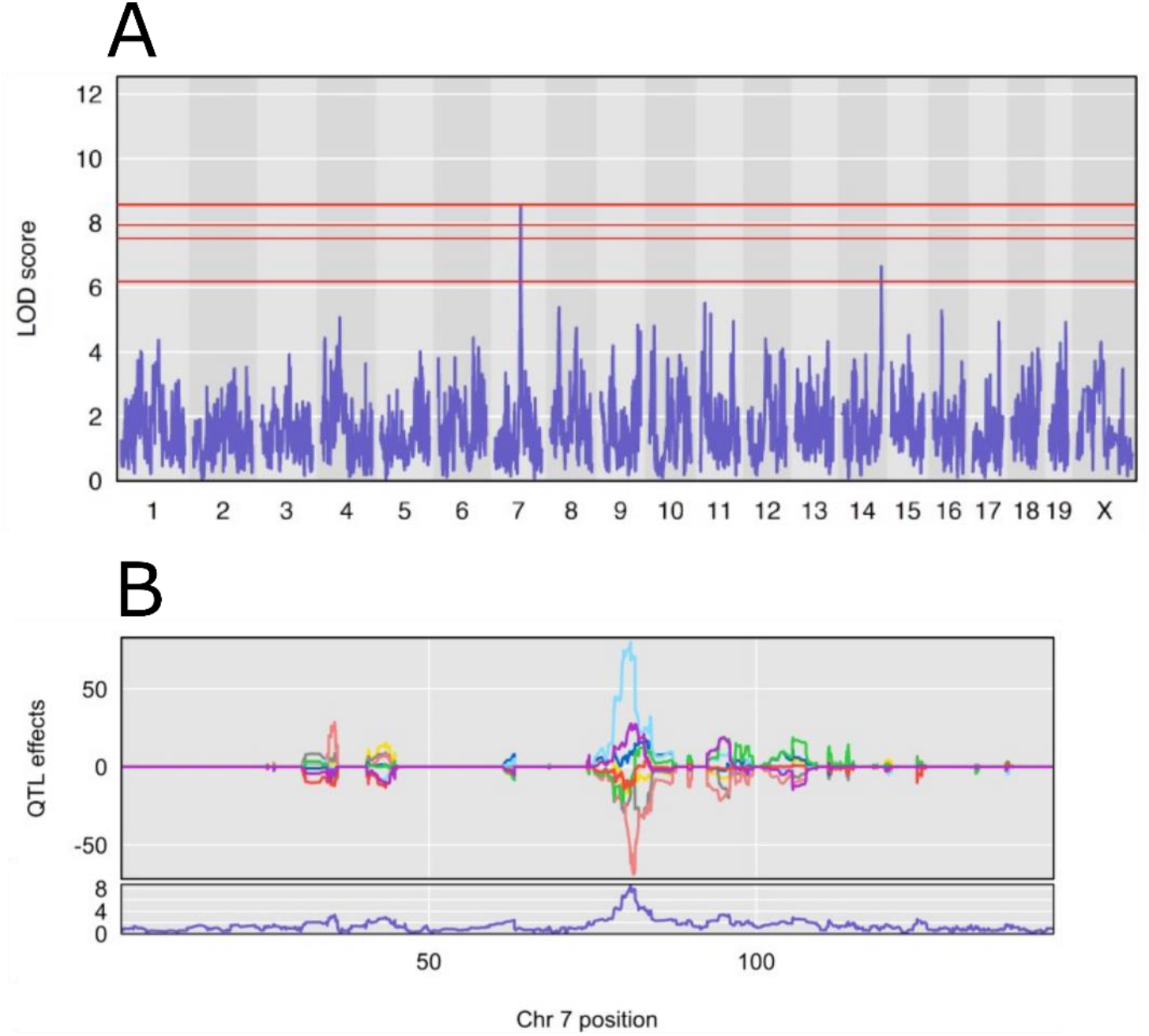
A significant QTL was mapped on chromosome 7 (80.26511-81.51397 Mb) for the reversal learning difference score, indicating one or more variants this locus associates with reversal learning. Haplotype analysis indicated the NZO/HlLtJ haplotype associated with larger difference scores and the 129/SvlmJ haplotype associated with smaller scores.

The QTL interval contained 58 genes. 24 of these genes were associated with *cis*-eQTL (Table 2). When these genes were assessed for strain-level correlation to reversal learning outcomes in 33 CC strains (Bailey et al. 2021), three were found to positively correlate with the reversal learning difference score (*2900076A07Rik, Wdr73 and Zscan2)*.

**Table 2.**
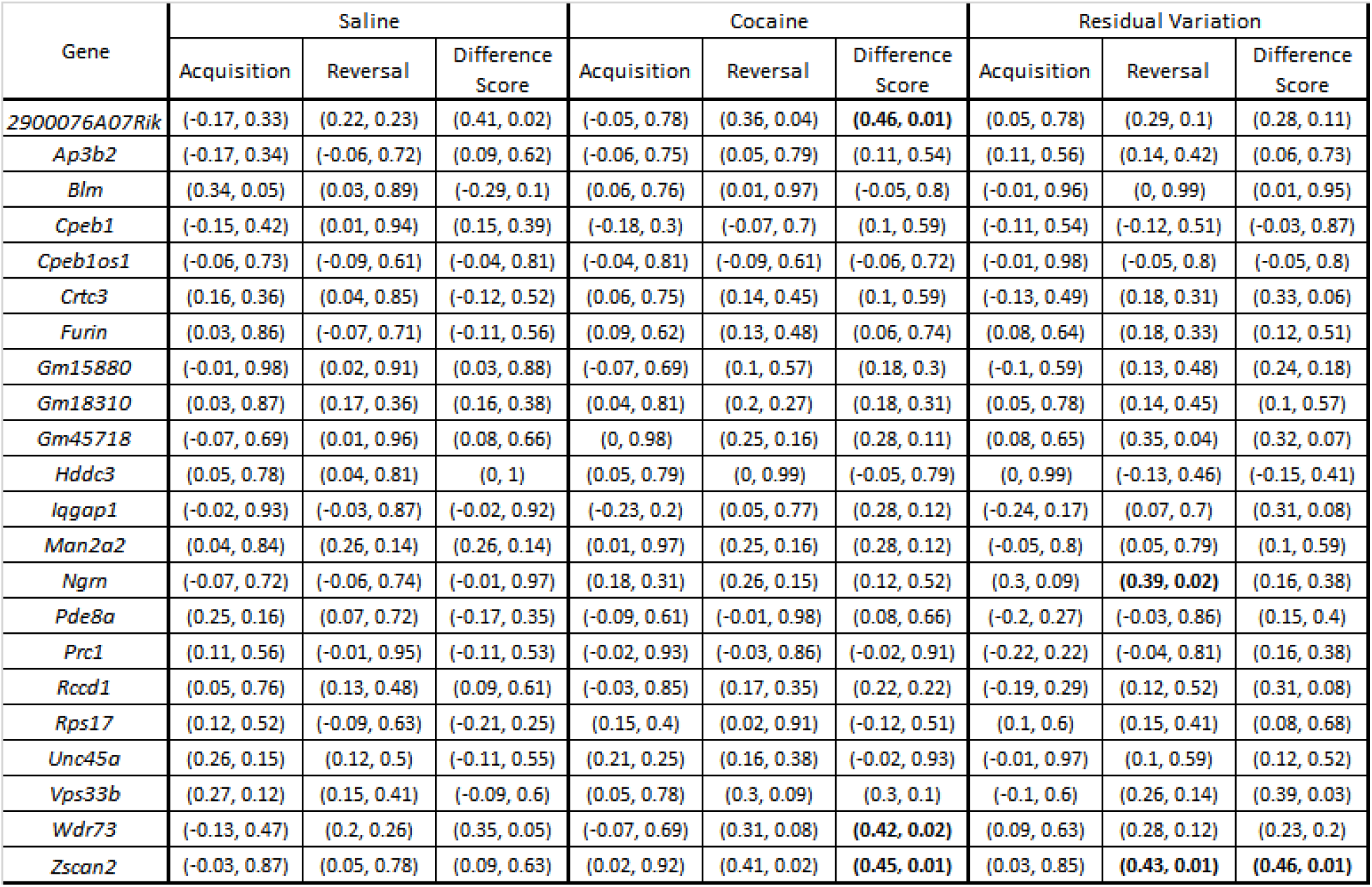
Correlation (r, p-value) between gene expression in 33 cocaine or saline exposed CC strains and reversal learning in independent groups of the same strains. Text in bold indicates a significant p-value.

Prioritized candidate genes were assessed by ePheWAS (systems-genetics.org) (Li et al., 2018) for correlation between BXD strain-level expression levels in the striatum or FC and all traits in the genenetwork.org phenome database. The candidate gene, *Wdr73,* demonstrated genetic correlations to dopamine receptor traits including: D1/D2 ratio (genenetwork ID 15554), D1 expression (genenetwork ID 15185), D2 expression (genenetwork ID 15186) and expression signature of D1 medium spiny neurons (genenetwork ID15552).

A suggestive QTL on chromosome 17 (position is in GRCm38 Mbp): Chr 17, Peak = 65.68404, LOD = 9.136811, Confidence Interval = 64.84549 - 66.34104) was detected for premature responses on the incorrect aperture in the reversal stage (Fig. 5A). The additive effects of haplotypes indicated the NZO/HILtJ haplotype associated with greater premature responding (Fig. 5B). The QTL interval contains 17 genes and 8 of these genes demonstrated *cis*-eQTL (Table 3). However, no genes demonstrated a correlation between gene expression and premature responses. Genes with *cis*-eQTL were also assessed for correlation to the reversal learning difference score. Expression of *Ralbp1* in the cocaine group demonstrated a positive correlation to the reversal learning difference score. Analysis by ePheWAS revealed that this gene is associated with acquisition of a visual discrimination operant response (genenetwork ID 16202) and aggregate protein formation on a Huntington’s disease model crossed to the BXD panel (genenetwork ID 16190). Furthermore, the *Ralbp1* gene harbors a non-synonymous variant (Table 4). Considering independent evidence that indicates *Ralbp1* may influence a similar operant task to that tested here, this gene may be considered an interesting candidate for further examination.

**Figure 5.**
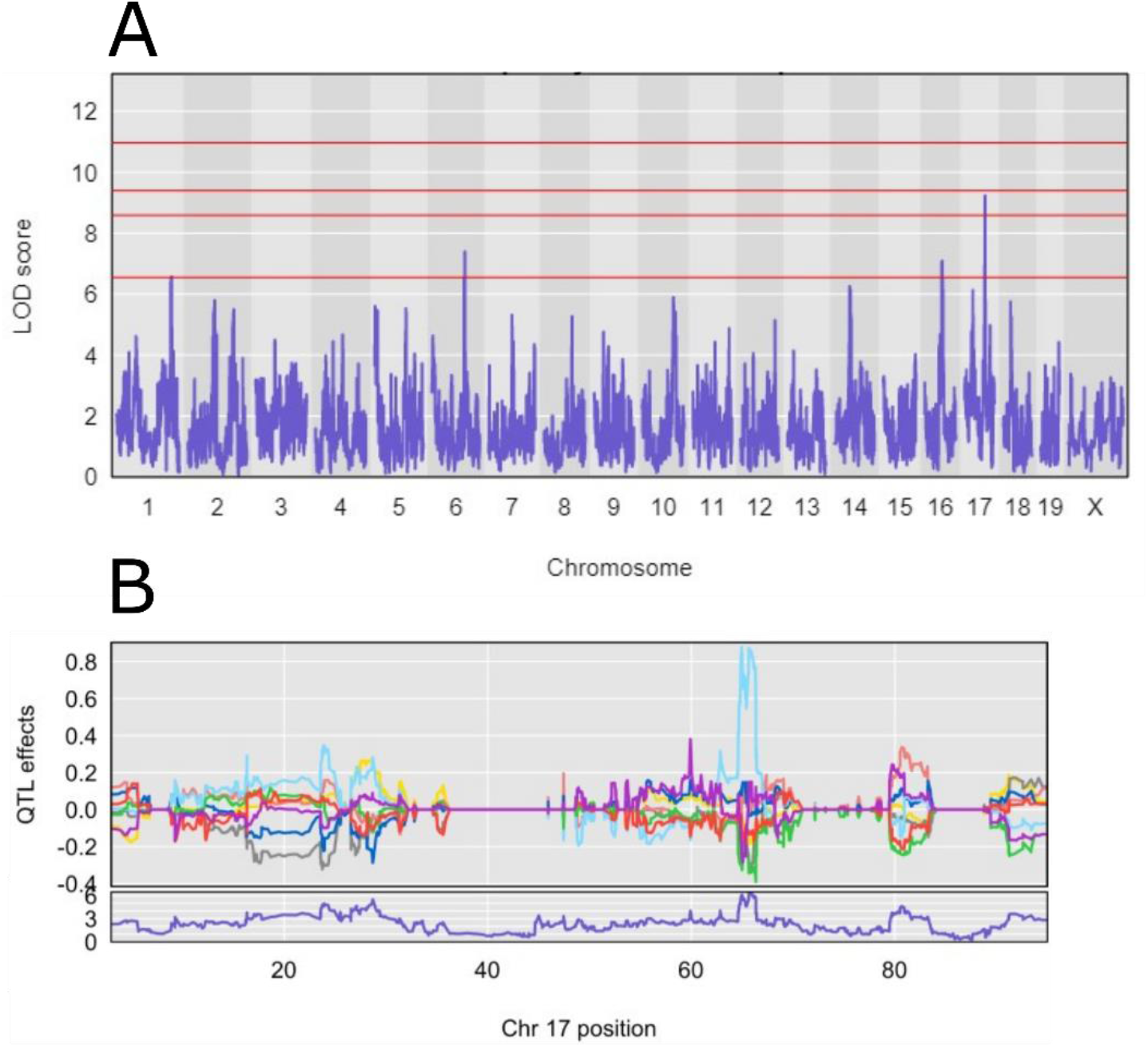
A suggestive QTL was mapped on chr 17 (64.84549 - 66.34104 Mb) for premature responding. Haplotype analysis indicated the NZO/HlLtJ associated with greater premature responding.

**Table 3.** Correlation (r, p-value) between gene expression in 33 cocaine or saline exposed CC strains and premature responding during reversal learning in independent groups of the same strains.

| Gene | eQTL.LOD | Sham | Coc | Res |
| --- | --- | --- | --- | --- |
| Ankrd12 | 26.12 | (0, 0.99) | (0.02, 0.93) | (0.02, 0.93) |
| Ddx11 | 19.63 | (-0.13, 0.47) | (-0.1, 0.6) | (0.01, 0.95) |
| Ppp4r1 | 14.68 | (0.1, 0.59) | (0.03, 0.87) | (-0.04, 0.82) |
| Rab31 | 9.41 | (0.31, 0.08) | (0.17, 0.36) | (0.05, 0.77) |
| Ralbp1 | 29.44 | (0.21, 0.25) | (0.17, 0.33) | (0.02, 0.9) |
| Twsg1 | 46.59 | (0.02, 0.92) | (0.13, 0.48) | (0.21, 0.23) |
| Vapa | 28.49 | (0.06, 0.76) | (0.07, 0.7) | (0.05, 0.8) |
| Washc1 | 15.22 | (0.07, 0.7) | (0.07, 0.69) | (0.03, 0.86) |

**Table 4.** Positional candidate genes (within the 1.5 lod interval) with genetic variants.

| Gene | Chr | Gene Type | Gene ID (MGI) | 3 prime UTR | Splice region | Downstream | Inter-genic | Intron | Intron, non coding | Non coding exon | Upstream gene | Synonymous | Mis-sense | Structural |
| --- | --- | --- | --- | --- | --- | --- | --- | --- | --- | --- | --- | --- | --- | --- |
| 1700023D08Rik | 7 | unclassified | 1921473 | 0 | 0 | 0 | 0 | 0 | 0 | 0 | 0 | 0 | 0 | DEL |
| Alpk3 | 7 | protein coding | 2151224 | 0 | 0 | 4 | 0 | 10 | 0 | 0 | 5 | 0 | 0 |  |
| Ap3b2 | 7 | protein coding | 1100869 | 0 | 0 | 0 | 0 | 0 | 1 | 0 | 1 | 0 | 0 |  |
| Cpeb1 | 7 | protein coding | 108442 | 0 | 0 | 18 | 0 | 126 | 11 | 0 | 1 | 0 | 0 |  |
| Crtc3 | 7 | protein coding | 1917711 | 0 | 0 | 1 | 0 | 0 | 0 | 0 | 0 | 0 | 0 |  |
| Gm15544 | 7 | pseudo | 3782993 | 0 | 0 | 0 | 0 | 0 | 0 | 0 | 1 | 0 | 0 |  |
| Gm18392 | 7 | pseudo | 5010577 | 0 | 0 | 0 | 0 | 0 | 0 | 0 | 0 | 0 | 0 | DEL |
| Gm18922 | 7 | pseudo | 5011107 | 0 | 0 | 0 | 0 | 0 | 0 | 0 | 0 | 0 | 0 | DEL |
| Gm32112 | 7 | lncRNA | 5591271 | 0 | 0 | 0 | 0 | 0 | 0 | 0 | 0 | 0 | 0 | DEL |
| Gm32178 | 7 | lncRNA | 5591337 | 0 | 0 | 0 | 0 | 0 | 0 | 0 | 0 | 0 | 0 | DEL |
| Gm42398 | 7 | lncRNA | 5625283 | 0 | 0 | 0 | 0 | 0 | 0 | 0 | 0 | 0 | 0 | DEL |
| Gm45991 | 7 | lncRNA | 5825628 | 0 | 0 | 0 | 0 | 0 | 0 | 0 | 0 | 0 | 0 | DEL |
| Iqgap1 | 7 | protein coding | 1352757 | 0 | 0 | 0 | 0 | 0 | 0 | 0 | 0 | 1 | 0 |  |
| Nmb | 7 | protein coding | 1915289 | 0 | 0 | 0 | 0 | 0 | 0 | 0 | 7 | 0 | 0 |  |
| Pde8a | 7 | protein coding | 1277116 | 0 | 0 | 5 | 0 | 74 | 23 | 0 | 10 | 0 | 0 | INS |
| Platr32 | 7 | lncRNA | 3801726 | 0 | 0 | 1 | 0 | 0 | 10 | 0 | 8 | 0 | 0 | INS |
| Ralbp1 | 7 | protein coding | 108466 | 0 | 0 | 0 | 0 | 0 | 0 | 0 | 0 | 0 | 1 |  |
| Rps17 | 7 | protein coding | 1309526 | 0 | 0 | 6 | 0 | 0 | 0 | 0 | 0 | 0 | 0 |  |
| Sec11a | 7 | protein coding | 1929464 | 0 | 0 | 4 | 0 | 0 | 21 | 3 | 6 | 0 | 0 | INV |
| Slc28a1 | 7 | protein coding | 3605073 | 1 | 1 | 4 | 0 | 9 | 0 | 0 | 0 | 0 | 0 | INS |
| Twsg1 | 7 | protein coding | 2137520 | 1 | 0 | 0 | 0 | 0 | 0 | 0 | 0 | 0 | 0 |  |
| Vapa | 7 | protein coding | 1353561 | 0 | 0 | 0 | 0 | 1 | 0 | 0 | 0 | 0 | 0 |  |
| Washc1 | 7 | protein coding | 1916017 | 0 | 0 | 0 | 0 | 0 | 0 | 0 | 0 | 1 | 0 |  |
| Wdr73 | 7 | protein coding | 1919218 | 0 | 0 | 6 | 0 | 0 | 10 | 0 | 12 | 0 | 0 | INV |
| Zfp592 | 7 | protein coding | 2443541 | 0 | 0 | 2 | 0 | 2 | 0 | 0 | 1 | 0 | 0 |  |
| Zscan2 | 7 | protein coding | 99176 | 0 | 0 | 38 | 0 | 14 | 0 | 0 | 8 | 0 | 0 | INS |
| Ankrd12 | 17 | protein coding | 1914357 | 0 | 0 | 1 | 0 | 0 | 0 | 0 | 1 | 0 | 0 |  |
| Ddx11 | 17 | protein coding | 2443590 | 0 | 0 | 0 | 0 | 4 | 0 | 0 | 0 | 0 | 0 |  |
| Gm23264 | 17 | snRNA | 5453041 | 0 | 0 | 0 | 0 | 0 | 0 | 0 | 1 | 0 | 0 |  |
| Mtcl1 | 17 | protein coding | 1915867 | 0 | 0 | 14 | 0 | 0 | 0 | 0 | 0 | 0 | 0 | INS |
| Ppp4r1 | 17 | protein coding | 1917601 | 0 | 0 | 0 | 0 | 0 | 3 | 0 | 0 | 0 | 0 |  |
| Rab31 | 17 | protein coding | 1914603 | 1 | 0 | 1 | 0 | 4 | 0 | 0 | 1 | 0 | 0 |  |
| Tmem232 | 17 | protein coding | 2685786 | 0 | 0 | 0 | 0 | 1 | 0 | 0 | 0 | 0 | 0 |  |
| Txndc2 | 17 | protein coding | 2389312 | 0 | 0 | 0 | 0 | 1 | 0 | 0 | 2 | 0 | 0 |  |

## Discussion

Impulsive action is a heritable trait that associates with risk for SUDs (Bailey et al., 2021; Brewer & Potenza, 2008; Calu et al., 2007; Camchong et al., 2011; Cervantes et al., 2013b; Dalley et al., 2011; de Wit, 2009; Gullo et al., 2010; Izquierdo & Jentsch, 2012; J. Jentsch, 2002; Perry & Carroll, 2008; Smith et al., 2015), and to some degree, this association may be due to a genetic correlation (coheritability). As a consequence, identifying the genetic regulators of impulsive behaviors may indirectly illuminate SUD genetics and neurobiology. We have previously found that the Collaborative Cross (CC) inbred strains and their founders demonstrate heritable variation in impulsive action, as measured by the reversal learning task (Bailey et al., 2021). In the present study, we utilized the Diversity Outbred (DO) mice, derived from the same founders as the CC strains, to characterize reversal learning and perform genome-wide QTL mapping to discover loci that may influence reversal learning traits. As expected, DO mice demonstrated a broad range of reversal learning performance. Our analyses of these data revealed a significant QTL that influenced reversal learning performance and a suggestive QTL that influenced premature responding.

The difference score for reversal learning captures the relative difficulty subjects have in adapting to the unexpected switch in response-outcome contingencies that happens at reversal. On average, trials to criterion are greater in the reversal stage, producing a positive difference score. however the range of performance in the DO mice is broad, with some mice taking ~200 fewer trials in reversal while mice at the other extreme required >300 additional trials to complete the reversal stage relative to acquisition. This variation is, in part, due to genetic differences in the DO mouse and is thus amenable to genome-wide QTL studies. QTL mapping revealed a significant QTL on chromosome 7 for this trait. The broadly defined confidence interval contained 58 genes. Gene expression data from the DO mice and 33 CC strains was utilized to determine positional candidate genes on the basis of striatum *cis*-eQTL and heritable expression patterns that are correlated with reversal learning difference scores in the same CC strains. This analysis indicated three genes as top candidates (*2900076A07Rik*, *Wdr73, Zscan2)*.

Further analysis of these prioritized genes by ePheWAS of publicly available gene expression and phenome datasets in the BXD recombinant inbred mouse panels revealed that *Wdr73* associated with heritable variation in striatal dopamine receptor transcript expression. Given the importance of striatal dopamine in reversal learning and risk for SUDs (Bergstrom et al., 2020; Clarke et al., 2008; Cools et al., 2009; Everitt & Robbins, 2013), *Wdr73* may impact reversal learning by affecting dopamine system function in this brain region. Furthermore, mutations in *Wdr73* are associated with Galloway-Mallowat syndrome, a developmental/neurological disorder (Rosti et al., 2016) and this gene was recently highlighted as a positional candidate in a multivariate GWAS of mood disorders and psychosis in human subjects (Mallard et al., 2019). Given the collection of evidence to suggest *Wdr73* may influence comorbid psychiatric conditions and striatal dopamine, this gene is considered a top candidate.

Premature responding during reversal learning is a measure of impulsive action analogous to measures in five choice serial reaction time (Bari et al., 2008). Given that this trait demonstrated a very modest correlation to the reversal learning differences score, it may provide unique and valuable genetic information. DO mice demonstrated a broad range of premature responding (near 0 to ~ 6 premature responses per trial). We discovered a suggestive QTL for premature responding on chromosome 17. The confidence interval contained 17 positional candidate genes. Eight of these genes have striatum cis-eQTL; however, none demonstrated genetic correlation to premature responding. These genes were also tested for genetic correlation to reversal learning difference scores. The gene *Ralbp1* positively correlated to differences scores, and ePheWAS analysis of his gene revealed that it is genetically correlated to phenotypes gathered in a similar operant discrimination task in the BXD mouse panel (genenetwork ID 16202). Additionally, this gene also correlated to aggregate protein formation in a Huntington’s disease model that was tested across BXD strains (genenetwork ID 16190). This gene also harbors a non-synonymous variant. Collectively, this evidence may indicate *Ralbp1* a candidate gene for further consideration.

The DO and CC mouse populations are genetically diverse mouse resources that have proven valuable for the study of impulsive action and addiction genetics. We have utilized the DO mice to follow up previous research in the CC strains that indicated reversal learning is heritable in these populations and amenable to forward genetic approaches. This approach has revealed a novel QTL for reversal learning difference scores and a suggestive QTL for premature responding during reversal learning. Additional work is underway to characterize cocaine self-administration and other traits related cocaine use disorder in the DO/CC populations (Kim et al., 2021; Saul et al., 2020; Schoenrock et al., 2020). Future analysis will integrate data presented here with these additional studies to facilitate further discovery of the genetics that simultaneously influence impulsivity and SUD-related traits.

## Acknowledgements

These studies were supported, in part, by Public Health Service grants P50-DA039841(EJC, JDJ, LGR, LMT), P30-CA034196 (Lutz, Cathleen M.; VMP) and T32-AA025606 (JDJ and JRB). The authors have no conflicts of interest to declare.

